# Sagittal Subtalar and Talocrural Joint Assessment Between Barefoot and Shod Walking: A Fluoroscopic Study

**DOI:** 10.1101/378216

**Authors:** Benjamin D. McHenry, Karen M. Kruger, Emily L. Exten, Sergey Tarima, Gerald Harris

## INTRODUCTION

Despite the fact that shoes are routinely worn during typical daily activities [1, 2], intra-foot kinematic models are usually developed and reported in the barefoot condition [3–5]. Understanding shod foot kinematics requires the description of foot/bone position inside footwear. This description has proven challenging using standard stereophotogrammetry techniques. Most external marker based studies in the current literature that report on shod foot mechanics use sandals [6–9], remove shoe material to expose the underlying anatomic area for marker placement [6, 10, 11], or place markers on the outer surface of the shoe [12–14]. While these approaches have increased understanding of shod foot mechanics, they have their limitations.

Sandals cannot be assumed to replicate shoes, as differences in foot biomechanics will occur due to the restriction of foot motion from upper shoe materials, lacing, and morphology [15]. A 2009 study by Hagen and Hennig reported significant differences in maximum pronation velocity, and peak pressure at various foot locations dependent on shoe lacing pattern while running [16]. Shoes with windowing (holes cut into them to expose anatomic locations for marker placement) have also been found to alter the mechanical properties of the shoe. A 2012 study by Shultz and Jenkyn determined that hole sizes larger than 1.7 x 2.5 cm would disrupt shoe integrity [17], and a 2006 study by Bulter et al. reported a 10% reduction in heel counter stability from shoe windowing [18]. In addition, attempts to measure intra-foot kinematics using external markers placed on the outer surface of a shoe present methodological concerns. A 2011 study by Bishop et al. reported static vector magnitudes up to 16.7 mm between markers placed on the external surface of the shoe and the underlying anatomic landmark [1]. These static discrepancies between marker placement and the underlying anatomy lead to dynamic kinematic differences as measured in a study by Reinschmidt et al. that reported tibiocalcaneal plantarflexion/dorsiflexion differences up to 8.1 degrees between shoe mounted and bone mounted markers.

Recently, fluoroscopy has been used to overcome these challenges in using external marker based foot models to measure shod foot kinematics. The radiographic nature of fluoroscopy allows for real time dynamic visualization of the underlying anatomy with no footwear modifications. Several fluoroscopic studies appear in the literature reporting on barefoot walking kinematics [19–23], but very few report on shod kinematics [24, 25]. In a 2014 study by Campbell et al., tibiocalcaneal kinematics between barefoot and shod walking were compared for six subjects, though talocrural (talus relative to tibia) and subtalar (calcaneus relative to talus) motion were not measured [24]. Wang et al. compared talocrural and subtalar kinematics from barefoot and shod walking in four cm and 10 cm high-heels, however only six individual images were analyzed between heel strike and toe-off [25].

The purpose of the current study was to determine sagittal plane talocrural and subtalar kinematic differences between barefoot and shod walking in athletic shoes. Dynamic fluoroscopic imaging was used to determine talocrural and subtalar kinematics from heel strike to 80% stance. The kinematic model used in this study applies a combination of motion and fluoroscopic data, and has previously been used to describe barefoot [20, 22] as well as Control Ankle Movement (CAM) boot walking [21].

## MATERIALS AND METHODS

In this Institutional Review Board (IRB) approved study, 13 male subjects (mean age 22.9 ± 2.9 years, mean weight 77.2 ± 6.9 kg, mean height 178.2 ± 3.7 cm), screened for exclusion criteria gave written informed consent prior to being tested while walking barefoot and in athletic walking shoes (New Balance Men’s MW927 Health Walking Shoe) (Figure 1). Exclusion criteria included any significant injury to the foot/ankle or any previous lower extremity surgery (bilateral).

**Figure 1.**
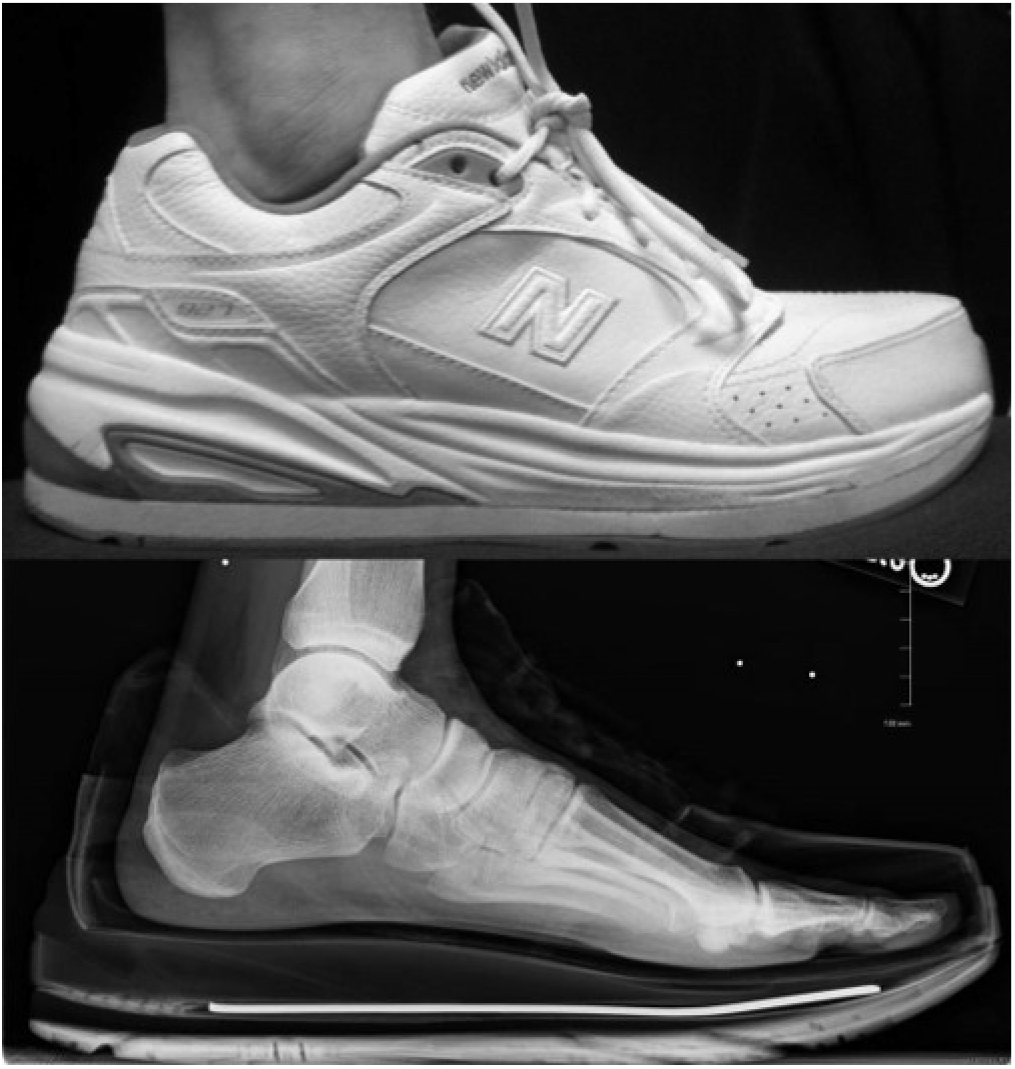
Athletic walking shoe (top). Static x-ray in athletic walking shoe (bottom).

The data capture system consisted of a modified fluoroscopy unit[20] placed within an existing Vicon motion analysis capture volume (Vicon Motion Systems, Inc., Oxford, UK). The fluoroscopy unit (OEC 9000, GE, Fairfield, CT) was modified so that the image intensifier and emitter could be set on opposite sides of the width of the walkway (Figure 2). Heel strike and toe-off events were detected using an embedded multi-axis force plate (AMTI, Watertown, MA).

**Figure 2.**
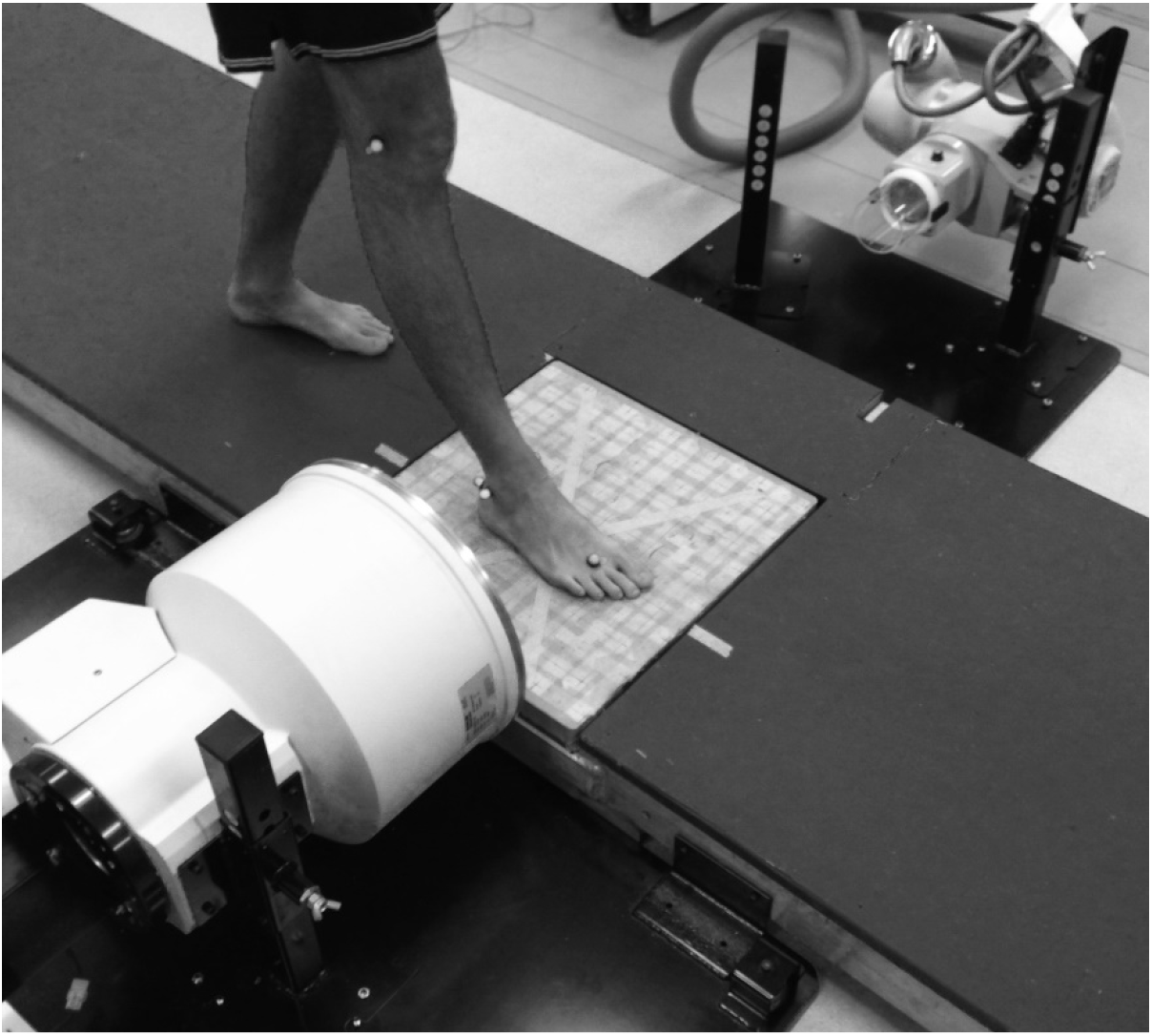
System configuration showing the walkway, emitter (far side), and image intensifier (nearside). Also shown is typical foot placement during image collection. Reprinted from McHenry et al. Foot Ankle Int. 2017 Nov;38(11):1260-1266.

The right leg and foot of each subject were instrumented with six reflective markers (d = 16 mm) in accordance with Table 1. For the shod trials, foot markers (calcaneal tuberosity and head of second metatarsal) were placed directly on the athletic walking shoe as close to their corresponding anatomic locations as possible. Fluoroscopic and motion data were collected simultaneously (120 Hz) as subjects walked along a custom walkway. Each subject completed five trials walking barefoot and five trials walking shod. Following dynamic data collection, static right foot x-rays were taken for each subject in both conditions (barefoot and shod) (Figure 1).

Radiation restrictions obviated recollection of fluoroscopic data if there was improper foot placement in the capture volume. Because of this, not all 13 subjects had five trials of data for each condition. For the barefoot condition, subjects averaged 4.5 ± 0.5 trials; for the athletic walking shoe condition, subjects averaged 3.8 ± 1.3 trials. Subject foot placement also determined how much of stance phase could be analyzed for each trial collected. If the tibia vacated the fluoroscopic field of view during toe-off, the analysis stopped. No portions of stance phase were analyzed for which fewer than ten subjects had data.

The kinematic model used in this study has been previously applied to describe barefoot talocrural and subtalar sagittal plane motion [20, 22]. The model uses external marker position to define a tibial local coordinate system, and fluoroscopic markers to define the talar and calcaneal local coordinate systems. External markers (medial/lateral malleoli and medial/lateral femoral epicondyles) were used to define the tibial local coordinate system as only the very distal end of the tibia was fluoroscopically visible for much of stance. For the talus and calcaneus, two points of interest per bone (talus, calcaneus) were translated from pixel coordinates to motion analysis global coordinates using a method of global referencing. This method has previously been shown to have errors less than 2 mm with subject foot progression angles of ± 5 degrees [20]. Average foot progression angle for the current study was 3.3 degrees external for barefoot and 4.8 degrees external for shod. These translated points of interest were defined in the sagittal plane of the foot and were then used to describe local coordinate systems for the talus and calcaneus. These local coordinate systems were used to calculate talocrural and subtalar sagittal plane kinematics, with motion defined as distal position relative to proximal. Kinematics were also calculated from the statically collected data, and these angles served as neutral position. Kinematic repeatability using this system has been determined to be 1.06 degrees [20].

A statistical analysis was done comparing shod to barefoot kinematics as well as temporal spatial parameters. The four sagittal plane kinematic positions statistically analyzed were talocrural plantarflexion at heel strike, talocrural peak plantar flexion, subtalar dorsiflexion at heel strike, and subtalar peak dorsiflexion. The three temporal spatial parameters statistically analyzed were walking speed, cadence, and stride length.

The Shapiro Wilk test was performed on each metric analyzed for testing the null hypothesis that differences between shod and barefoot were normally distributed. For metrics that were normally distributed, a paired t-test was performed with the null hypothesis that the true mean difference between shod and barefoot walking was zero. Wilcoxon signed rank test was planned as a nonparametric alternative to the paired t-test. Statistical significance was declared at p ≤ 0.05.

## RESULTS

All kinematic trials were averaged within a subject and then among subjects for assessment of each tested condition (barefoot and shod). Talocrural kinematics for both conditions are shown in Figure 3. At heel strike, barefoot plantarflexion was 4.2 ± 4.8 degrees and shod was −2.4 ± 5.3 degrees. Barefoot peak plantarflexion was 10.9 ± 4.3 degrees occurring at 11% stance, and shod peak plantarflexion was 7.0 ± 4.7 degrees occurring at 16% stance.

**Figure 3.**
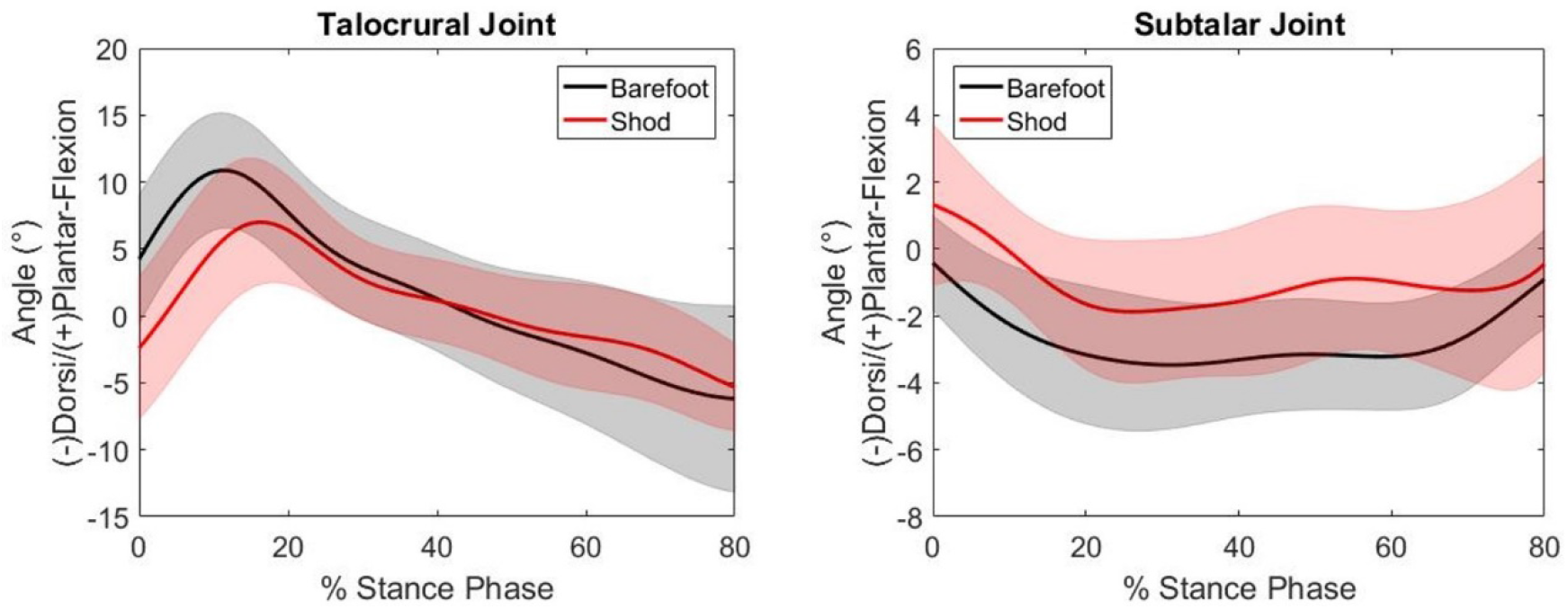
Talocrural (left) and Subtalar (right) plantar/dorsiflexion angles during stance. The black solid line and grey band represents the mean and standard deviation of all thirteen subjects walking barefoot. The red solid line and light red band represents the mean and standard deviation of all thirteen subjects walking in the athletic walking shoes.

Subtalar kinematics for both conditions are shown in Figure 3. At heel strike, barefoot dorsiflexion was −0.4 ±1.4 degrees and shod was 1.3 ± 2.4 degrees. Barefoot peak dorsiflexion was −3.5 ± 1.9 degrees occurring at 31% stance, and shod peak dorsiflexion was 1.9 ± 2.1 degrees occurring at 26% stance.

Using the Shapiro Wilks normality test, all metrics analyzed were normally distributed. Using a paired t-test it can be seen that all metrics analyzed were statistically significant between shod and barefoot, with the exception of subtalar dorsiflexion at heel strike and subtalar peak dorsiflexion (Table 2).

## DISCUSSION

As a result of wearing shoes, the current study shows average walking speed significantly increased by 0.04 m/s, average stride length significantly increased by 0.10 m, and average cadence significantly decreased by 4.74 steps/min (Table 2). Similar results have been reported in other studies comparing barefoot to shod motion [11, 26–29]. In a 120 subject study of children (6-13 years old), Moreno-Hernandez et al. reported an increase in walking speed of 0.0537 m/s, an increase of stride length of 0.0733 m, and a decrease in cadence of 3.51 steps/min (all statistically significant) [28]. In a study of 980 children (5-27 years old) Lythgo et al. reported an increase in walking speed of 0.08 m/s, an increase of stride length of 0.111 m, and a decrease in cadence of 3.9 steps/min (all statistically significant)[26]. The current study temporal spatial results compare favorably to these previous studies and confirm that the natural response to footwear is an increase in walking speed and stride length, with a reduction in cadence.

Comparing barefoot to shod talocrural joint kinematics (Table 2), wearing shoes significantly reduced plantarflexion at heel strike (−6.73 degrees). In addition, wearing shoes significantly reduced talocrural peak plantarflexion during loading response (−3.22 degrees). It can also be seen that wearing shoes delayed peak talocrural joint plantarflexion during loading response (Figure 3). Barefoot peak plantarflexion occurred at 11% stance, while shod peak plantarflexion occurred at 16% stance. This effect of shoes causing decreased ankle joint plantarflexion at heel strike [6, 11, 14, 24, 29], decreased peak plantarflexion during loading response [6, 14, 24, 29], and a delay in peak plantarflexion during loading response [6, 11, 14, 29] has been previously reported. The current study, however, is the first to attribute this trend directly to talar motion relative to tibia.

Comparing barefoot to shod subtalar joint kinematics (Table 2), wearing shoes increased dorsiflexion at heel strike (1.80 degrees). In addition, wearing shoes increased subtalar peak dorsiflexion during loading response (1.18 degrees). This increased subtalar joint dorsiflexion was sustained throughout stance phase. Between heel strike and 80% stance, shod subtalar joint dorsiflexion was, on average, 2.0 degrees less dorsiflexed than barefoot. The only other study to directly measure subtalar joint motion during shod walking was done by Wang et al. in 2016 which used low-heeled (four cm) and high-heeled (10 cm) shoes [25]. This makes a direct comparison to the current study difficult. While the Wang study showed no significant plantar/dorsiflexion position differences comparing barefoot to low-heels, they did report significantly less dorsiflexed subtalar positions comparing barefoot to high-heels [25]. While the current study results reflect this trend of shod subtalar dorsiflexion being less than barefoot, the paired t-test showed no statistical significance.

The current study used fluoroscopic technology to compare shod walking with barefoot. The results showed that walking while wearing shoes significantly increased average walking speed and stride length while significantly reducing average cadence. At the talocrural joint, walking in shoes significantly reduced plantarflexion at heel strike and significantly reduced peak plantarflexion. At the subtalar joint, wearing shoes while walking reduced dorsiflexion during the entire portion of stance phase analyzed (heel strike to 80%). The current study is limited in that it used a single fluoroscope with a focused analysis on the sagittal plane [20]. To report on coronal or transverse motion, the fluoroscope could be repositioned relative to the walkway. A biplane fluoroscopic system would be required to report on simultaneous triaxial kinematics, and our group has recently published the technical details of such a system [30]. While a second fluoroscope would allow for the measurement of triaxial motion, it would increase radiation exposure, which may not be necessary for all studies. While the dose was low, use of ionizing radiation was required for the current work. This estimated radiation exposure was ten μSv/trial, well below the United States Nuclear Regulatory Commission (USNRC) whole body annual occupational limits of five rems (50,000 μSv).

## Acknowledgments

The contents of this article were developed under a grant from the National Institute on Disability, Independent Living, and Rehabilitation Research (NIDILRR grant number 90AR5022-01-00 Formerly H133P140023-14). NIDILRR is a Center within the Administration for Community Living (ACL), Department of Health and Human Services (HHS). The contents of this article do not necessarily represent the policy of NIDILRR, ACL, HHS, and you should not assume endorsement by the Federal Government.

